# Transcriptome analysis in whole blood reveals increased microbial diversity in schizophrenia

**DOI:** 10.1101/057570

**Authors:** Loes M Olde Loohuis, Serghei Mangul, Anil PS Ori, Guillaume Jospin, David Koslicki, Harry Taegyun Yang, Timothy Wu, Marco P Boks, Catherine Lomen-Hoerth, Martina Wiedau-Pazos, Rita M Cantor, Willem M de Vos, René S Kahn, Eleazar Eskin, Roel A Ophoff

## Abstract

The role of the human microbiome in health and disease is increasingly appreciated. We studied the composition of microbial communities present in blood across 192 individuals, including healthy controls and patients with three disorders affecting the brain: schizophrenia, amyotrophic lateral sclerosis and bipolar disorder. By using high quality unmapped RNA sequencing reads as candidate microbial reads, we performed profiling of microbial transcripts detected in whole blood. We were able to detect a wide range of bacterial and archaeal phyla in blood. Interestingly, we observed an increased microbial diversity in schizophrenia patients compared to the three other groups. We replicated this finding in an independent schizophrenia case-control cohort. This increased diversity is inversely correlated with estimated cell abundance of a subpopulation of CD8+ memory T cells in healthy controls, supporting a link between microbial products found in blood, immunity and schizophrenia.

## Introduction

Microbial communities in and on the human body represent a complex mixture of eukaryotes, bacteria, archaea and viruses. In recent years, mounting evidence has demonstrated the involvement of the microbiome in human health and disease. In particular, through the ‘microbiota-gut-brain-axis’ (1, 2), the microbiome has been implicated in complex psychiatric disorders, including schizophrenia and major depressive disorder(3-8), possibly via an impact on intestinal permeability(9).

High-throughput sequencing offers a powerful culture-independent approach to study the underlying diversity of microbial communities in their natural habitats across different human tissues (10) and diseases (3, 11-15). The majority of current microbiome studies use fecal samples and target 16S ribosomal RNA gene sequencing (16). With the availability of comprehensive compendia of reference microbial genomes and phylogenetic marker genes (17), it has become feasible to use non-targeted sequencing data to identify the microbial species across different human tissues and diseases in a relatively inexpensive and easy way.

Other than in cases of sepsis, we currently lack a comprehensive understanding of the human microbiome in blood, as blood has been generally considered a sterile environment lacking proliferating microbes (18). However, over the past few decades, this assumption has been challenged (19, 20), and the presence of a microbiome in the blood has received increasing attention (21-23).

To explore potential connections between the microbiome and diseases of the brain, we performed a comprehensive analysis of microbial products detected in blood in almost two hundred individuals, including patients with schizophrenia, bipolar disorder and sporadic amyotrophic lateral sclerosis. These three disease groups represent complex polygenic traits that affect the central nervous system with largely unknown etiology. Moreover, roles for the microbiome in all the diseases have been previously hypothesized (5, 24-26). We used available high quality RNA sequencing (RNA-Seq) reads from whole blood that fail to map to the human genome as candidate microbial reads for microbial classification. We observed an increased diversity of microbial communities in schizophrenia patients, and we replicated this finding in an independent dataset. Careful analyses, including the use of positive and negative control datasets, suggest that these detected phyla represent true microbial communities in whole blood and are not present in samples due to contaminants. With the increasing number of RNA-Seq data sets, our approach may have great potential for application across different tissues and disease types.

## Materials and Methods

A brief description of Materials and Methods follows below; see Supplementary Methods for the full details.

### Sample Description

The discovery sample consists of unaffected controls (Controls, n=49) and patients with three brain-related disorders: schizophrenia (SCZ, n=48), amyotrophic lateral sclerosis (ALS, n=47) and bipolar disorder (BPD, n=48). The replication sample includes Controls (n=88) and SCZ samples (n=91). Sample recruitment of the cohorts is described in the Supplementary Methods. All study methods were approved by the institutional review board of the University of California at Los Angeles, San Francisco or the Medical Research Ethics Committee of the University Medical Center Utrecht at The Netherlands. All participants provided written informed consent.

### Sample sequencing

For the discovery sample, RNA-Seq libraries were prepared using Illumina’s TruSeq RNA v2 protocol, including ribo-depletion protocol (Ribo-Zero Gold). In total, we obtained 6.8 billion 2x100bp paired-end reads for the primary study (35.3M ± 6.0 paired-end reads per sample). The replication sample was processed at the same core facility using the same standardized procedures as the discovery sample. However, the RNA-Seq libraries were prepared with poly(A) enrichment, a procedure more selective than the total RNA that was used for the discovery sample. A total of 3.8 billion reads were obtained (26.3M ± 12.0).

### Sequence Analysis

We separated human and non-human reads, and use the latter as candidate microbial reads for taxonomic profiling of microbial communities. To identify potentially microbial reads, we developed the following pipeline. First, we filtered read pairs and singleton reads mapped to the human genome or transcriptome. Because total number of reads may affect microbial profiling, we performed normalization by sub-sampling to 100,000 reads for each sample. Next, we filtered out low-quality and low-complexity reads using FASTX and SEQCLEAN (see urls). Finally, the remaining reads were realigned to the human references using the Megablast aligner (27) in order to exclude any potentially human reads. The remaining reads were used as candidate microbial reads in subsequent analyses. Figure 1 displays an overview of our pipeline.

**Figure 1.** Microbial profiling using RNA-Seq data from whole blood. (A) We analyzed a cohort of 192 individuals from four subject groups, i.e. Schizophrenia (SCZ, n=48), amyotrophic lateral sclerosis (ALS n=47), bipolar disorder (BPD n=48), unaffected control subjects (Controls n=49). (B) Peripheral blood was collected for RNA collection. (B) RNA-Seq libraries were prepared from total RNA using ribo-depletion protocol. Reads that failed to map to the human reference genome and transcriptome were sub-sampled and further filtered to exclude low-quality, low complexity, and remaining potentially human reads. High quality, unique, non-host reads were used to determine the taxonomic composition and diversity of the detected microbiome. See also Table S1.

### Taxonomic profiling

To access the assembly and richness of the microbiomial RNA in blood, we used phylogenetic marker genes to assign the candidate microbial reads to the bacterial and archaeal taxa. We used PhyloSift (v 1.0.1 with default parameters) to perform taxonomic profiling of the whole blood samples (17). PhyloSift makes use of a set of protein coding genes found to be relatively universal (i.e., present in nearly all bacterial and archaeal taxa) and have low variation in copy number between taxa. Homologs of these genes in new sequence data (e.g., the transcriptomes used here) are identified and then placed into a phylogenetic and taxonomic context by comparison to references from sequenced genomes. For our replication study, we used MetaPhlAn for microbial profiling v.1.7.7(28). MetaPhlAn was run in 2 stages; the first stage identifies the candidate microbial reads (i.e., reads hitting a marker) and the second stage profiles meta-genomes in terms of relative abundances. We used MetaPhlAn, rather than PhyloSift, due to differences in library preparation (polyA enrichment versus Ribo-Zero); there were an insufficient number of reads matching the database of the marker genes curated by PhyloSift for adequate microbial profiling of the replication sample.

### Estimating Microbial diversity

Microbial diversity, or alpha diversity, within each sample was determined using the inverse Simpson index. This index simultaneously assesses both richness (corresponding to the number of distinct taxa) and relative abundance of the microbial communities within each sample (29). In particular, it enables effective differentiation between the microbial communities shaped by the dominant taxa and the communities with many taxa with even abundances (30) *(asbio* R package). To measure sample-to-sample dissimilarities between microbial communities, we use Bray-Curtis beta diversity index, which accounts for both changes in the abundances of the shared taxa and for taxa uniquely present in one of the samples *(vegan* R package). Higher beta diversity indicates higher level of dissimilarity between microbial communities, providing a link between diversity at local scales (alpha diversity) and the diversity corresponding to total microbial richness of the subject group (gamma diversity (31)).

### Statistical analysis of microbiome diversity

To test for differences in alpha diversity between disease groups, we fit an analysis of covariance (ANCOVA) model using normalized values of alpha, including sex and age, and technical covariates (RNA INtegrity value (RIN), batch, flow cell lane and RNA concentration) into the model. Bonferroni correction for multiple testing was used. To determine the relative effect size of alpha diversity on schizophrenia status, we fit a logistic regression model including the same covariates and measure reduction in R^2^ comparing the full logistic regression model versus a reduced model with alpha removed. Analysis of beta diversity was performed analogously (see Supplementary Methods).

### Reference-free microbiome analysis

We complement the reference-based taxonomic analysis with a reference independent analysis. We use EMDeBruijn (https://github.com/dkoslicki/EMDeBruijn), a reference-free approach capable of quantifying differences in microbiome composition between the samples. EMDeBruijn compresses the k-mer counts of two given samples onto de Bruijn graphs and then measures the minimal cost of transforming one of these graphs into the other. To determine overlap between the results from PhyloSift and EMdeBruin, we correlated principal components of EMdeBruin and PhyloSift by Spearman rank correlation, including all samples.

### Estimation of cell proportions in whole blood

We assessed DNA methylation data from 65 controls taken from our replication sample, and we compared methylation-derived blood cell proportions estimated using Houseman’s estimation method (32, 33) to alpha diversity after adjusting for age, gender, RIN and all technical parameters. We tested whether alpha diversity levels are associated with cell type abundance estimates. More details on the method, quality control pipeline of the methylation data and statistical analysis can be found in Supplementary Methods.

## Results

### Studying microbial RNA in blood

To study the composition of microbial RNA in blood, we determined the microbial meta-transcriptome present in the blood of unaffected controls (Controls, n=49) and patients with three brain-related disorders: schizophrenia (SCZ, n=48), amyotrophic lateral sclerosis (ALS, n=47) and bipolar disorder (BPD, n=48) (Figure 1, Table 1).

Using our filtering pipeline, an average of 33,546 of 100,000 unmapped reads are identified as high quality, unique non-host reads and were used as candidate microbial reads in our analyses. From these, PhyloSift was able to assign an average of 1,235 reads (1.24% ± 0.41%, mean ± standard deviation) to the bacterial and archaeal gene families. A total of 1,880 taxa were assigned, with 23 taxa at the phylum level (Figure 2). Most of the taxa we observed derived from bacteria (relative genomic abundance 89.8% ± 7.4%), and a smaller portion derived from archaea (relative genomic abundance 12.28% ±6.4%).

**Figure 2.** Relative abundances of microbial taxa at phylum level. Phylogenetic classification is performed using PhyloSift, which is able to assign the filtered candidate microbial reads to the microbial genes from 23 distinct taxa on the phylum level.

In total, we observed 23 distinct microbial phyla with on average 4.1 ± 2.0 phyla per individual. The large majority of taxa observed in our sample is not universally present in all individuals; the single exception is Proteobacteria, which dominates all samples with 73.4% ± 18.3% relative abundance (Figure 2 dark green color). Several bacterial phyla show a broad prevalence across individuals and disorders (present in 1/4 of the samples of each subject group). Those phyla include Proteobacteria, Firmicutes and Cyanobacteria, with relative abundance 73.4% ± 18.3%, 14.9 ±10.9%, and 11.0% ± 8.9% (Table S2). This is in line with recent published work on the blood microbiome using 16S targeted metagenomic sequencing reporting relative abundance of 80.4-87.4% and 3.0-6.4% for Proteobacteria and Firmicutes, respectively (23). The other two phyla identified in this study (Actinobacteria and Bacteroidetes) were also detected in our sample in more than 25 individuals. Although Proteobacteria and Firmicutes, are commonly associated with the human microbiome (34), some members of these phyla might be associated with reagent and environmental contaminants (35, 36).

To validate our pipeline and investigate the possibility of contamination introduced during RNA isolation, library preparation and sequencing steps, we performed both negative and positive control experiments (see Supplementary Results and Methods for details). In brief: no microbiome sequences were detected in transcriptome data in lymphopblast cell lines (negative control), and we only detected the Chlamydiae phylum in RNA-Seq from cells infected with Chlamydiae (positive control). We examined experimental procedures and technical parameters on microbial composition, and we observed no link between the presence of microbial communities and possible confounders.

To compare the inferred microbial composition found in blood with that in other body sites, we used taxonomic composition of 499 meta-genomic samples from Human Microbiome Project (HMP) obtained by MetaPhlAn or five major body habitats (gut, oral, airways, and skin) (10). Of the 23 phyla discovered in our sample, 15 were also found in HMP samples, of which 13 are confirmed by at least ten samples. Our data suggest that the predominant phyla detected in blood are most closely related to the known oral and gut microbiome (Table S2). Comparing the microbial composition of whole blood with the microbiome detected in atherosclerotic plaques (37), we observe that the four phyla that together make up for >97% of the microbiome in plaques are also identified in our sample (Firmicutes, Bacteroidetes, Proteobacteria, Actinobacteria).

Finally, it should be noted that the sequencing technology does not allow for identification of the origin of microbial RNA. That is, we can not distinguish whether the observed microbial signatures in blood are originate from bacterial communities actually present in the blood, or whether the RNA crossed into the blood stream from elsewhere.

### Increased microbial diversity in schizophrenia samples

To evaluate potential differences in microbial profiles of individuals with the different disorders (SCZ, BPD, ALS) and unaffected controls, we explored the composition and richness of the microbial communities across the groups.

We observed increased alpha diversity in schizophrenia samples compared to all other groups *(ANCOVA* P < 0.005 for all groups, Figure 3a, Table 2 and Table S5, *Bonferroni* correction). These differences are corrected for covariates and are independent of potential confounders, such as experimenter and RNA extraction run (Figure S1 and S2), and they are not the consequence of a different number of reads being detected as microbial in schizophrenia samples (see Supplementary Results). No significant differences were observed between the three remaining groups (BPD, ALS, Controls). In our sample, alpha diversity was found to be a significant predictor of schizophrenia status and explained 5.0% of the variation as measured by reduction in Nagelkerke’s R^2^ from logistic regression. We observe no correlation between polygenic risk scores (39) and alpha diversity in our schizophrenia sample (n= 32, Kendall’s tau= 0.008, *P* = 0.96, Supplementary Methods). We also did not observe differences in alpha diversity between sexes or across ages, nor are our results driven by the relatively younger schizophrenia cohort (Supplementary Results). Alpha diversity at other main taxonomic ranks yields a similar pattern of increased diversity in schizophrenia (Figure S3).

**Figure 3.** Increased diversity of microbiome detected in blood in schizophrenia samples. (A) Alpha diversity per sample for four subject groups (Controls, ALS, BPD, SCZ), measured using the inverse Simpson index on the phylum level of classification. Schizophrenia samples show increased diversity compared to all three other groups (ANCOVA P < 0.005 for all groups, after adjustment of covariates, see also Methods, Table S5 and Figure S3). (B) Alpha diversity per sample of schizophrenia cases and controls, measured using the inverse Simpson index on the genus level of classification. Schizophrenia samples show increased within-subject diversity compared to Controls (P = 0.003 after adjustment of covariates).

The increased diversity observed in schizophrenia patients may be due to specific phyla characteristic to schizophrenia, or due to a more general increased microbial diversity in people affected by the disease. To investigate this, we compared diversity across individuals within the schizophrenia group to control samples. We compared beta diversity across pairs of samples with schizophrenia and controls, resulting in three subject groups: SCZ_Controls, SCZ_SCZ and Controls_Controls. The lowest diversity was observed in the Controls_Controls group (0.43 ± 0. 21), followed by SCZ_SCZ (0.50 ± 0.14), and the highest beta diversity values were observed for SCZ_Controls (0.51 ± 0.17) (P< 0.05 for each comparison, by ANCOVA after correcting for three tests). This result was confirmed by permanova (P<0.001) based on 1,000 permutations. Thus, the observed increased alpha diversity in schizophrenia is not caused by a particular microbial profile, but most likely represents a non-specific overall increased microbial burden (see also Figure S4 and Supplementary Results).

In addition to measuring individual microbial diversity (alpha), and diversity between individuals (beta), we measured the total richness of the microbiome by the total number of distinct taxa of the microbiome community observed within an entire subject group (gamma diversity (40)). We observed that all 23 distinct phyla are observed in schizophrenia: gamma(SCZ)=23 compared to gamma(Controls)=20, gamma(ALS)=16 and gamma(BPD)=18.

We complemented reference-based methods (PhyloSift and MetaPhlAn) with EMDeBruijn, a reference-independent method. EMDeBruin distances measured between samples correlated significantly with beta diversity (Spearman rank P < 2.2e-16, rho = 0.37, including SCZ and Controls). Also, EMDeBruijn PCs correlated with principal components obtained from edge PCA based on the PhyloSift taxonomic classification (correlation between EMDeBruijn PC1, and PhyloSift PC1 is P = 1.824e-09; Spearman rank correlation is rho = -0.42; see also Figure S5). After correcting covariates, the first three EMDeBruijn PCs are significant predictors of schizophrenia status, and jointly explained 7.1% of the variance (P< 0.05 for each PC).

### Group differences of individual phyla

In addition to a global difference between schizophrenia and the other groups, we also investigated whether there are particular individual phyla contributing to the differences between schizophrenia and other groups. There are two phyla detected more often in schizophrenia cases versus all the other groups: Plactomycetes, observed in 20 SCZ cases compared to 3(ALS) 2(BPD) 5(Controls) (P= 0.0002 Fisher’s exact for four groups, Bonferroni corrected for 23 tests P=0.0057) and Thermotogae, observed in 20 SCZ cases compared to 6 ALS, 3 BPD and 6 Controls (P= 0.0006 Fisher’s exact, corrected P=0.014). No outliers were observed for the other groups (see Table S7).

### Replication

We performed a replication experiment in an independent case-control sample: schizophrenia (SCZ n=91) and healthy controls (Controls n=88) (See Table S1.D). MetaPhlAn was able to assign 5,174 reads (0.089% ± 0.039%, mean ± standard deviation) on average to the bacterial gene families.

Schizophrenia samples showed increased alpha diversity on genus level (2.73 ± 0.77 for cases, versus 2.32 ± 0.57 for controls, corrected P = 0.003 Figure 3b) and explained 2.5% of variance as measured by reduction in Nagelkerke R^2^, thus replicating our main finding of increased diversity in schizophrenia. While our original analysis was performed on the phylum level, in our discovery sample we observe a similar increase of diversity at the genus level (see Figure S3). Similar to our discovery cohort, we observed no significant correlation between alpha diversity and age or differences across gender. Beta diversity and EMDeBruijn analyses also show similar, though not identical, patterns of nonspecific increased diversity in schizophrenia samples (Supplementary Results).

### Cell type composition and diversity

We hypothesized that differences in microbial diversity may be linked to whole blood cell type composition. Our analysis shows that the proportion of one cell type, CD8^+^ CD28^-^ CD45RA^-^ cells, is significantly negatively correlated with alpha diversity after correction for all other cell-count estimates as estimated from whole blood DNA methylation data (correlation = -0.41, P=7.3e-4, n= 65 Controls from the Replication study, Figure S6, Table S6). These cells are T cells that lack CD8^+^ naïve cell markers CD28 and CD45RA and are thought to represent a subpopulation of CD8^+^ memory T cells (41, 42). We observed that low alpha diversity correlates with high levels of cell abundance of this population of T cells.

## Discussion

We used high throughput RNA sequencing from whole blood to perform microbiome profiling and identified an increased diversity in schizophrenia patients.

While other studies of human microbiome using RNA-Seq have been conducted (43, 44), this is the first assessing the microbiome from whole blood by using unmapped non-human reads. Despite the fact that transcripts are present at much lower fractions than human reads, we were able to detect microbial transcripts from bacteria and archaea in almost all samples. The microbes found in blood are thought to be originating from the gut as well as oral cavities (45, 46), which is in line with our finding that the microbial profiles found in our study most closely resemble the gut and oral microbiome as profiled by the HMP (10). The taxonomic profile of the cohort samples suggests the prevalence of the several phyla, Proteobacteria, Firmicutes and Cyanobacteria, across individuals and different disorders included in our study. This is in line with a recent study that used 16S targeted meta-genomic sequencing, which reported Proteobacteria and Firmicutes among the most abundant phyla detected in blood (23).

Rigorous quality control is critically important for any high-throughput sequencing project, especially in the context of studying the microbiome (35). To this end, we performed both negative and positive quality control experiments, and we carefully evaluated possible contamination effects introduced during the experiments. Our results suggest that the detected phyla represent true microbial communities in whole blood and are not due to contaminants. However, it should be noted that whether only the microbial products crossed into the bloodstream or whether the microbes themselves are present in blood cannot be answered using sequencing techniques. Future experiments, for example, using microscopy, culturing or direct measures of gut permeability, may be able to shed light on this question.

The most striking finding of our study that relates to diseases affecting the central nervous system is the increased microbial alpha diversity in schizophrenia patients compared to controls and the other two disease groups (ALS, BPD). We replicate this finding in an independent cohort of schizophrenia cases and controls. The replication experiment, while based on different library preparation (Ribo-Zero versus Poly(A)), provides strong evidence for an increased alpha diversity of the microbiome detected in blood in schizophrenia and explains roughly 5% of disease variation. We not only observe an increased individual microbial diversity, but also an increased diversity between individuals (Beta diversity) with schizophrenia compared to controls, rendering it unlikely that a single phylum or microbial profile is causing the disease-specific increase in diversity. Nevertheless, in our study we observed that two phyla in particular, Planctomycetes and Thermotogae, were present in significantly more schizophrenia samples when compared to the other groups. Interestingly, Planctomycetes is group of gram-negative bacteria closely related to Verrucomicrobia and Chlamydiae; together these comprise the Planctomycetes-Verrucomicrobia-Chlamydiae superphylum (47). From peripheral blood, infection with Chlamydiaceae species has been reported to be increased in schizophrenia (40%) compared to controls (7%) (48). Since Chlamydiae is one of the taxa of the superphylum, it is possible that the increase in Planctomycetes we observe is related to the observed increase in Chlamydiaceae species. As the collection of available reference genomes continues to grow and improve, future studies are needed to corroborate and refine these findings.

For the study of microbiome diversity, we employed reference-based methods (PhyloSift and MethPhlAn) and the EMDebruin method, a purely reference-agnostic approach. The latter showed strong correspondence to both reference-based methods, highlighting the value of this unbiased sequence-based analysis for investigating microbial differences across groups. However, in addition to differences in distribution of microbial transcripts, EMDebruin may capture variation of other, yet unknown, origin.

In addition to our observation that microbial diversity is more generally increased in schizophrenia, our study demonstrates the value of analyzing non-human reads present in the RNA-Seq data to study the microbial composition of a tissue of interest (49, 50). The RNA-Seq approach avoids biases introduced by primers in targeted 16S ribosomal RNA gene profiling. In addition, since *mRNA stability is low in prokaryotes,* RNA-Seq might offer a potential advantage of avoiding contamination of genomic DNA by dead cells compared to genome sequencing (51). Given the many large-scale RNA-Seq datasets that are becoming available, we anticipate that high-throughput meta-transcriptome-based microbiome profiling will find broad applications as a hypothesis-generating tool in studies across different tissues and disease types.

The increased microbial diversity observed in schizophrenia could be part of the disease etiology (i.e., causing schizophrenia) or may be a secondary effect of disease status. In our sample, we observed no correlation between increased microbial diversity and genetic risk for schizophrenia as measured by polygenic risk scores (52). In addition, it is remarkable that bipolar disorder, which is genetically and clinically correlated to schizophrenia (53), does not show a similar increased diversity. We did observe a strong inverse correlation between increased diversity and estimated cell abundance of a population of T-cells in healthy controls. Even though this finding is based on indirect cell count measures using DNA methylation data (41), the significant correlation highlights a likely close connection between the immune system and the blood microbiome, a relationship that has been documented before (54). More extensive cell count measures and/or better markers of immune sensing of microbial products could be used to study this relationship more directly. In the absence of a direct link with genetic susceptibility and the reported correlation with the immune system, we hypothesize that the observed effect in schizophrenia may be mostly a consequence of disease. This may be affected by lifestyle and/or health status differences of schizophrenia patients, including smoking, treatment plans, (chronic) infection, GI status, the use of probiotics, antibiotics and other drug use or other environmental exposures. Future targeted and/or longitudinal studies with larger sample sizes, detailed clinical phenotypes and more in-depth sequencing are needed to corroborate this hypothesis. Another interesting direction for future work is to study gut permeability in the context of our findings more directly. For example, how does damage to the gut (such as measured using I-FABP) affect observed microbial diversity in blood? These studies would likely result in an expanded understanding of the functional mechanisms underlying the connection between the human immune system, microbiome, and disease etiology. In particular, we hope that these future efforts will provide a useful quantitative and qualitative assessment of the microbiome and its role across the gut-blood barrier in the context of psychiatric disorders.

### Availability of Data and Materials

The data discussed in this publication have been deposited in NCBI’s Gene Expression Omnibus (55) and are accessible through GEO Series accession number GSE80974 (https://www.ncbi.nlm.nih.gov/geo/query/acc.cgi?acc=GSE80974).

## Acknowledgments

This work is supported by NIH/NIMH R01 5R01MH090553 (BP samples), 5R01NS058980 (ALS samples) and R01MH078075 (SCZ, Control samples), R21MH098035 (replication sample) awarded to R.A.O. L.M.O.L was financially supported by the National Institute of Neurological Disorders And Stroke of the National Institutes of Health under Award Number T32NS048004. The authors thank Dr. Jonathan Eisen for helpful discussions and insights throughout the course of this project. We thank Dr. Lana Martin for helpful edits to the manuscript. Finally, we thank the study subjects for their participation.

## Financial Declarations

The authors have no conflicts of interest to declare.

